# A dry method for preserving tear protein samples

**DOI:** 10.1101/131060

**Authors:** Weiwei Qin, Chan Zhao, Linpei Zhang, Ting Wang, Youhe Gao

## Abstract

Tears covering the ocular surface is an important bio-fluid containing thousands of molecules, including proteins, lipids, metabolites, nucleic acids, and electrolytes. Tears are valuable resources for biomarker research of ocular and even systemic diseases. For application in biomarker studies, tear samples should ideally be stored using a simple, low-cost, and efficient method along with the patient’s medical records. For this purpose, we developed a novel Schirmer’s strip-based dry method that allows for storage of tear samples in vacuum bags at room temperature. Using this method, tear protein patterns can also be preserved. Liquid chromatography-mass spectrometry/mass spectrometry analysis of proteins recovered from the dry method and traditional wet method showed no significant difference. Some tissue/organ enriched proteins were identified in tear, thus tear might be a good window for monitoring the change of these tissues or organs. This dry method facilitates sample transportation and enables the storage of tear samples on a large scale, increasing the availability of samples for studying disease biomarkers in tears.

## Introduction

Tears overlay the epithelial cells of the cornea and conjunctiva surface. It provides lubrication, protection, and nutrition to the ocular surface. Tear is a complex extracellular fluid, with normal human tears consisting of 1543 proteins [1], approximately 100 different types of small molecule metabolites [1, 2], and more than 600 lipid species from 17 major lipid classes [3]. Tear fluid can be easily and noninvasively accessed [4] and has become a useful resource for biomarker research of ocular and systemic diseases. According to a recent review [5], hundreds of potential specific molecular biomarkers in tears were found to be associated with ocular diseases such as dry eye disease, keratoconus, and Graves’ orbitopathy. Other reports showed that tears can also reflect the states of breast cancer, prostate cancer, and multiple sclerosis [5-7]. For example, studying early-stage biomarkers of breast cancer in tears, Lebrech et al. reported significant differences in tear proteins between breast cancer patients and healthy, showing 90 % specificity and sensitivity[8]. It is also reported a panel of 20 biomarkers with an overall specificity and sensitivity of 70 %[9]. Böhm et al. reported a distinctive difference in 20 biomarkers of breast cancer, versus healthy controls[10]. Moreover, tears may reflect central metabolism in some neurological disorders [6].

Tears show promise as biofluids for biomarker studies and should be preserved along with a patient’s medical record. This is a critical step in validation, which facilitates biomarker research and its translation from the bench to the bed. The primary methods for collecting tears are using the Schirmer’s strip and glass capillary tube, followed by flash-freezing at −80°C [11]. Cryopreservation of tears cannot absolutely prevent the degradation of proteins, as the samples contain various enzymes and hydrolases. Additionally, use of the required cold chain during sample transportation is challenging and costly.

Here, we dried the Schirmer’s strip soaked with tears and stored the strip in a vacuum bag. Importantly, the proteins were dry, preventing their degradation and enabling preservation at room temperature.

## Materials & Methods

### 1. Ethical statement

The consent procedure and this study protocol were approved by the Institutional Review Board of the Institute of Basic Medical Sciences, Chinese Academy of Medical Sciences. (Project No. 007–2014). Written informed consent was obtained from each subject.

### 2. Tear collection and preservation

Tear samples were collected from seven healthy volunteers by Schirmer’s type I tear test without using local anesthesia. No volunteers had a recent history of ocular disease or contact lens usage. The Schirmer’s strips were inserted for 5 min in the lower eyelid in standard fashion in both eyes by the same subject. Schirmer’s strips from both eyes of the volunteer were cut longitudinally into two halves immediately after sampling. Half of the sample was added to a 2-mL Eppendorf tube snap-frozen at −80°C for 2 weeks (Fig. 1A). The other half of the sample was stored by the dry method for 2 weeks (Fig. 1B). The strip soaked with tears was dried using a hair dryer (Philips HP8200) for 2–3 min, and then the strip was placed in an aseptic plastic bag. The bag was then sealed using a kitchen vacuum sealer and stored at room temperature.

**Figure 1.**
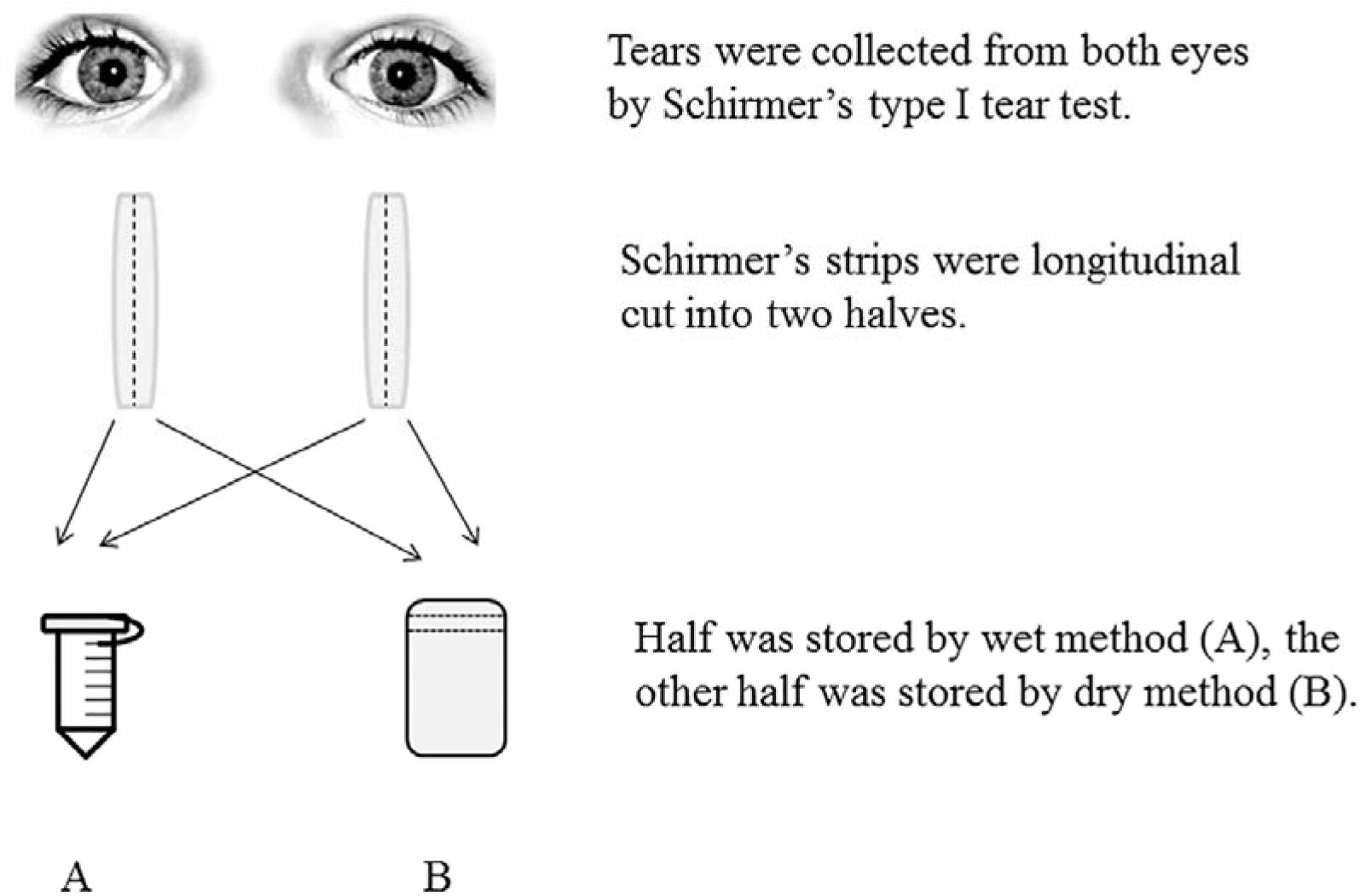
Workflow of the collection and preservation of tear samples. Wet method (A): strips were placed in a 2-mL Eppendorf tube snap-frozen at −80°C for 2 weeks; Dry method (B): strips were dried and then stored in a vacuum bag at room temperature for 2 weeks.

### 3. Protein extraction

The strip was cut into small pieces and transferred into a 0.6-mL tube. Next, 200 μL elution buffer (100 mM NH_3_HCO_3_, 50 mM NaCl) was added and gently shaken for 2 h at room temperature. The tube was punctured at the bottom with a cannula, placed in a larger tube (1.5 mL), and centrifuged at 12,000 g for 5 min [12]. The filtrate in the outer tube was collected and quantified by the Bradford method before SDS-PAGE analysis.

### 4. Tryptic digestion

Tear proteins were digested by filter-aided sample preparation methods [13]. Briefly, 200 μg of protein were loaded on the 10kD filter unit (Pall, USA), and 200 μl UA (8 M urea in 0.1 M Tris–HCl, pH 8.5) was added to the filter unit and centrifuged at 12,000g for 40 min. Then 200 μl ABC (0.05 M NH4HCO3 in water) was added and the centrifuged. Dithiothreitol solution (4.5 mM dithiothreitol in ABC) was added to the filter unit and incubated for 1 hour at 37°C. Centrifuge the filter units at 12,000g for 30 min. Iodoacetamide solution (10 mM iodoacetamide in ABC) was added to filter unit and incubated in the dark for 30 min at room temperature. Centrifuge the filter units at 12,000g for 30 min. Then the concentrate was dissolved in 50 mM NH4HCO3. Proteins were digested with trypsin (4 μg) for 14 h at 37°C. The digested peptides were desalted using Oasis HLB cartridges (Waters, USA). The resulting peptides were desalted and dried by a SpeedVac (Thermo Fisher Scientific, Waltham, MA, USA). The reproducibility of digestion was estimated, and the details was included in the supporting information.

### 5. LC-MS/MS analysis

The digested peptides were dissolved in 0.1% formic acid and loaded on a trap column (75 μm × 2 cm, 3 μm, C18, 100 Å). The eluent was transferred to a reversed-phase analytical column (50 μm × 150 mm, 2 μm, C18, 100 Å) by an Thermo EASY-nLC 1200 HPLC system. Peptides were analyzed using a Fusion Lumos mass spectrometer (Thermo Fisher Scientific). The Fusion Lumos was operated on data-dependent acquisition mode. Survey mass spectrometry (MS) scans were acquired in the Orbitrap using a 350–1550 m/z range with the resolution set to 120,000. The most intense ions per survey scan (top speed mode) were selected for collision-induced dissociation fragmentation, and the resulting fragments were analyzed in Orbitrap. Dynamic exclusion was employed with a 30-s window. Three technical replicate analyses were performed for each sample.

### 6. Data analysis

The MS/MS spectra were processed with Mascot software, using the human proteome database (UniProtKB/Swiss-Prot release 2014-01-10). The FASTA file contained 20120 protein sequences. Search parameters were set as follows: 10 ppm precursor mass tolerance, 0.02 Da fragment mass tolerance, two missed cleavage sites allowed in the trypsin digestion, cysteine carbamidomethylation as fixed modification, and oxidation (M) as variable modifications. Protein identifications were accepted if they could be established at greater than 91.0% probability [14] to achieve a false-discovery rate of less than 1.0% and contained at least 1 identified peptide.

## Results and Discussion

### 1. SDS-PAGE analysis of proteins recovered from tears stored by wet and dry methods

To estimate the effectiveness of storing tear proteins in a dry state at room temperature, proteins were recovered from strips that had been stored for 2 weeks by the wet method and dry method, and then separated by SDS-PAGE and stained with silver. As shown in Fig. 2, the same tear sample stored by the wet and dry methods exhibited similar legible patterns, suggesting that these two methods have similar effectiveness for protein preservation.

**Figure 2.**
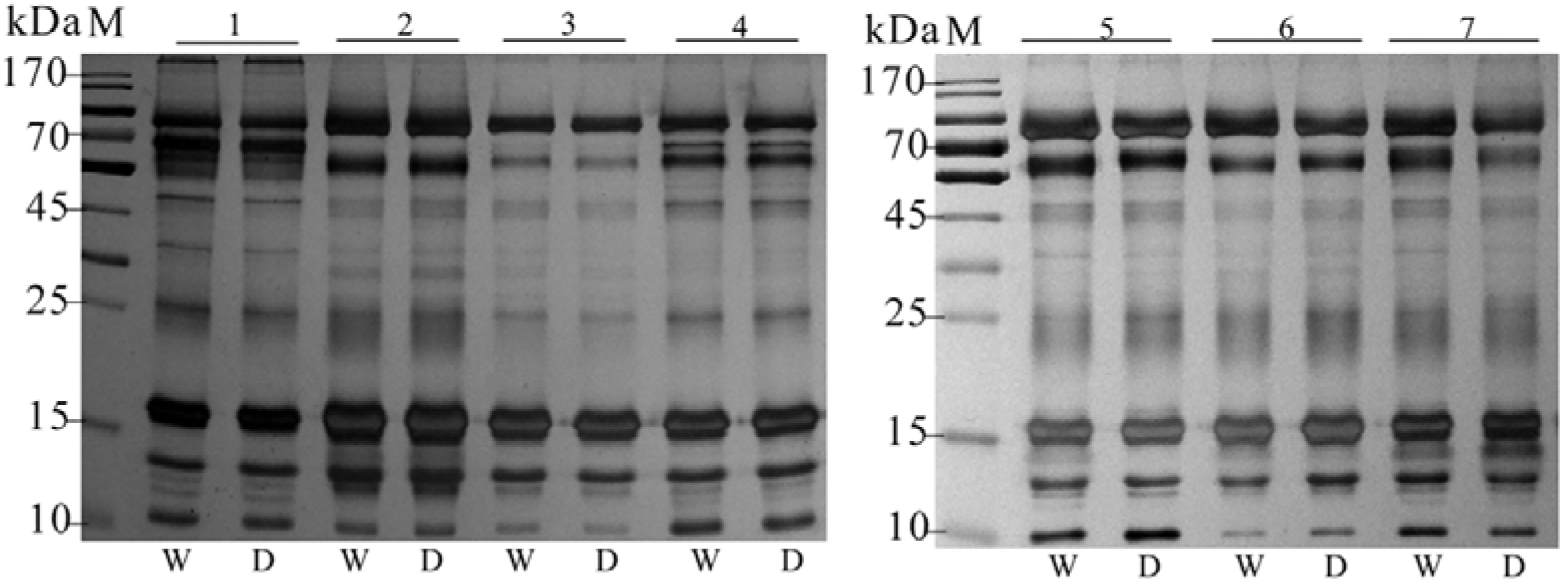
SDS-PAGE analysis of tear proteins recovered from wet method and dry method. In each lane, 5 μg tear proteins were loaded, stained with silver. M: marker; 1, 2, 3, 4, 5, 6, 7: different volunteers; W: wet method (strips soaked with tears was snap-frozen at −80°C); D: dry method (strips soaked with tears was dried and stored in a vacuum bag at room temperature).

### 2. LC-MS/MS identification of proteins recovered from wet method and dry method

To further evaluate the preservation effectiveness of the dry method, the exact species of proteins in the tear samples stored by the wet and dry methods, from 4 subjects, were identified by liquid chromatography (LC)-MS/MS. After label-free quantification, identification of proteins for three LC-MS/MS technical replicates prepared by the wet method revealed 316 shared proteins (456, 434, and 417 in each replicate), with an overlap rate of 72.5% (overlap rate = shared proteins/mean proteins × 100%) (Fig. 3A). Simultaneously, 290 shared proteins were identified (397, 394, and 395 in each replicate) within the replicates prepared by the dry method, with an overlap rate of 73.4% (Fig. 3B). A total of 240 proteins were identified as common by these two methods, with an overlap rate of 79.2% (Fig. 3C), which was similar to that of the LC-MS/MS technical replicates (Fig. S1, Fig. S2 and Fig. S3). Following preservation, some proteins showed differences between the wet method and dry method; thus, in each study, the same collection and preservation method should be used.

**Figure 3.**
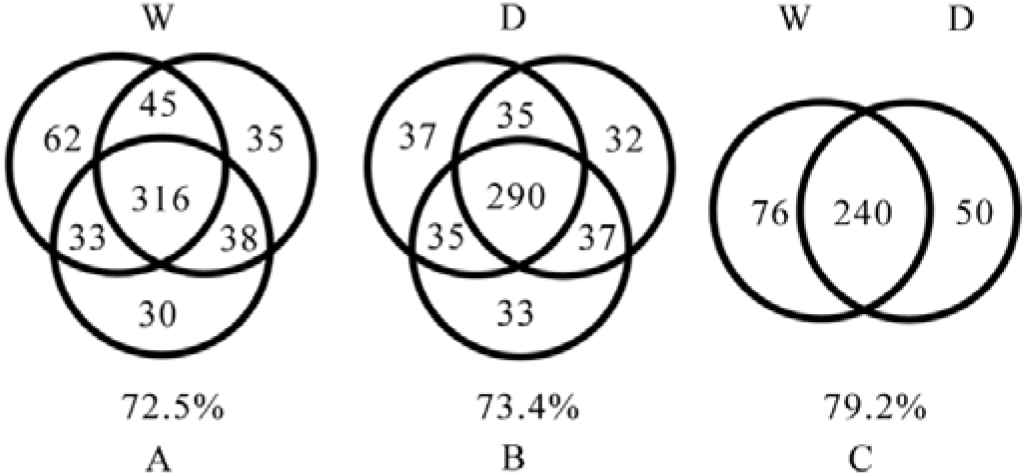
LC-MS/MS identification of tear samples from volunteer 1 preserved by two methods. A, B: Venn diagrams comparing proteins identified among 3 independent analyses of the sample preserved by the wet method (A) and dry method (B). C: Venn diagram comparing proteins identified between the wet and dry methods.

To compare further these two methods, spectral counting method was used to estimate each identified protein’s abundance [15]. We draw a correlation curve of identified proteins abundance between the dry and the wet method. The correlation coefficient (R2) of protein abundance was 0.9987, 0.9974, 0.9969, and 0.9993 respectively for volunteer 1, 2, 3 and 4 (Fig. 4, Fig. S4, Fig. S5 and Fig. S6). The results showed good correlation of protein abundance between these two methods. The protein only preserved in the wet or dry method were low abundance proteins.

**Figure 4.**
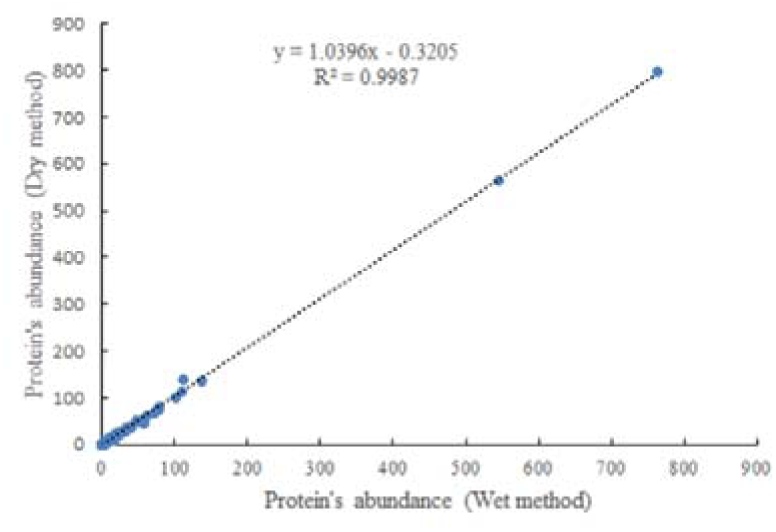
Correlation curve of identified proteins abundance between dry method and wet method for volunteer 1. Spectral counting method was used to estimate each identified protein’s abundance.

## Discussion

Recently, Uhlén et al reported a tissue-based map of the human proteome, describing the expression and distribution of human proteins across 44 different tissues and organs, both at the mRNA (32 tissues) and protein level [16]. In our study, we identified 514 tear proteins, and then we compared each of them to the tissue-enriched proteome. In all, 365 proteins that highly enriched in different tissues and organs were also identified in tears, and 132 proteins corresponding to 132 protein-encoding genes that highly expressed in different tissues and organs were also detected in tears (Table S1). This is an observation that tissue/organ enriched proteins are present in tear. There is no known mechanism as far as we know. At this point, we can only propose the possibility that if those organs had functional and/or structural changes, proteins enriched in those organs may be released in a different quality and quantity into the blood. These changes may somehow reach in tear and be reflected on proteins in tear. Therefore, tear might be a good window for monitoring the change of these tissues or organs. These proteins are not specific to those organs. They may also be made by tear gland locally.

According to the qualitative and quantitative result of LC-MS/MS, proteins only preserved in the wet or dry method were low abundance proteins. This is very likely caused by the proteomics strategy adopted in this study. LC-MS/MS with data dependent acquisition (DDA) was used, and it is based on signal intensity used for the precursor-ion selection, which result in an incomplete sampling of the peptide mixture generated to represent the proteome. The analytical reproducibility of peptide identifications obtained using DDA-based methods is about 75% overlap between technical replicates[17]. So we suggestion that most of the protein were preserved in both method. Even if there are differences, it would not be a problem when the same method is used in one study as we suggested.

In addition to proteins, other biomolecules in tears were preserved on the strip by the dry method, including lipids, metabolites, nucleic acids, and electrolytes. Since all tears were soaked on the strip, only water and some volatile matter was lost during the drying procedure. In this study, we only analyzed the proteins, considering the time consumption and experimental techniques. Seven volunteers were included. Because we focused on estimating the stability of this new dry method. This procedure including tear collection with Schirmer’s strip, protein preservation on dry and vacuum station, protein elution from the strip, and the protein identification using SDS-PAGE and LC-MS/MS. Except the preservation part, the other three parts have been proved to be reproducible respectively [1, 12, 13, 17]. To prove this dry method is stable, we studied all four parts’ reproducibility as a whole. Seven samples means seven technical replicates, it maybe not large enough for a biomarker study, but when considering that every step had been proved reproducible, it should be enough to prove the stability of the whole method.

We first reported the dry method for preserving tear samples. Compared to the wet method, the most significant difference between these two methods is on preservation procedure. By the wet method (the primary method), after the tear collection, the Schirmer’s strip is flash-freezing at −80°C. The disadvantage is that cryopreservation of tear samples cannot absolutely prevent the degradation of proteins, as the samples contain various enzymes and hydrolases. Additionally, use of the required cold chain during sample transportation is challenging and costly. By the dry method, after the tear collection, the Schirmer’s strip is dried and stored in a vacuum bag. The advantage is that the proteins were dry, preventing their degradation and enabling preservation at room temperature. Therefore, higher dryness degree and vacuum degree should keep tear samples at room temperature for longer period.

For all this, the dry method is applicable for establishing libraries of stored tear samples for long period and can simplify studies of disease biomarkers in tears.

## Acknowledgements

This work was supported by the National Key Research and Development Program of China (grant number 2016YFC1306300), the National Basic Research Program of China (grant number 2013CB530850), and funds from Beijing Normal University (grant numbers 11100704, 10300-310421102). The authors have declared no conflict of interest.

## Supporting Information

The raw data, the detail of the identified proteins have been uploaded to the data repository as supporting information, and all these data can be accessed on <u>https://figshare.com/s/b06a5a4e9f77b38ae6e1</u>.

